# Meta-analysis: a quantitative model of adult neurogenesis in the rodent hippocampus

**DOI:** 10.1101/2024.04.09.588728

**Authors:** Jon I. Arellano, Pasko Rakic

## Abstract

Contrary to humans, adult neurogenesis in rodents is not controversial. And in the last three decades, multiple studies in rodents have deemed adult neurogenesis essential for most hippocampal functions. The functional relevance of new neurons relies on their distinct physiological properties during their maturation before they become indistinguishable from mature granule cells. Most functional studies have used very young animals with robust neurogenesis. However, this trait declines dramatically with age, questioning its functional relevance in aging animals, a caveat that has been mentioned repeatedly, but rarely analyzed quantitatively. In this meta-analysis, we use data from published studies to determine the critical functional window of new neurons and to model their numbers across age in both mice and rats. Our model shows that new neurons with distinct functional profile represent about 3% of the total granule cells in young adult 3-month-old rodents, and their number decline following a power function to reach less than 1% in middle aged animals and less than 0.5% in old mice and rats. These low ratios pose an important logical and computational caveat to the proposed essential role of new neurons in the dentate gyrus, particularly in middle aged and old animals, a factor that needs to be adequately addressed when defining the relevance of adult neurogenesis in hippocampal function.

## Introduction

Adult hippocampal neurogenesis (AHN) has been a prolific topic of research and discussion for the last 30 years. The possibility of neuron renewal and the underlying promise of regeneration in the context of aging and neurological disease has been an important catalyst for the field that have attracted the attention of researchers, funding agencies, scientific journals, and the public. Due to the obvious limitations to perform studies on the human brain, functional inferences about adult neurogenesis have been collected almost exclusively in rodents, mostly in mice (Snyder, 2019). These studies include evaluating the effect of environmental or pharmacological interventions on AHN or using transgenic models of neurological diseases to study their effect on AHN (Snyder and Drew, 2020). Transgenic animals have also been used to study the effect of increased or decreased AHN on specific functions such as spatial memory, pattern separation or associative learning (e.g.: (Clelland et al., 2009; Drew et al., 2010; Arruda-Carvalho et al., 2011; Sahay et al., 2011). As a result, a large number and variety of functions have been related to adult neurogenesis, including: memory consolidation, flexible learning and updating, reward learning, emotional contextualization, time stamping, spatial contextualization and navigation, behavioral pattern separation, orthogonalization and avoidance of catastrophic interference, detection and seeking of novelty, regulation of affective behaviors and mood, and forgetting (Snyder and Drew, 2020; Kempermann, 2022).

Three factors must be considered when evaluating the functional contribution of adult born granule cells (CGs). The first is whether new neurons possess different connectivity or physiological responses that can confer them distinct functionality from that observed in mature neurons. The second, if their numbers at any given time are sufficient to have a functional impact in the circuit. And third, if their connectivity large enough and mature enough to support such functional contribution. Obviously, these conditions need to be met before any discussion about the possible function of new neurons can take place. While there is evidence supporting the first condition, there is less support for the other two. Interestingly, despite the many reviews on every possible aspect of AHN, those specific topics, namely if there are enough new neurons and if they are sufficiently connected have not received much attention (Snyder and Cameron, 2012). Due to space constrains, in this article we will address the first two questions, namely if new neurons have different physiology than mature cells and if they are numerous enough to support a relevant functional role in the adult rodent dentate gyrus, while the third question regarding the connectivity of new neurons will be analyzed in a separate article.

To answer the first two questions, we analyze available data on the pattern of connections and the physiology and maturation of adult born neurons in the dentate gyrus and we describe a model of quantitative neurogenesis that predicts the number of new neurons with distinct physiology that are available in the rodent dentate gyrus across age. The results show relatively low levels of functionally distinct new neurons at any age that decline quickly with age to reach very low levels in middle-aged and old animals. We compare our results with other quantitative analyses, and we discuss the implications of our findings in the context of hippocampal function.

<u>When is a rodent adult?</u>

After more than a century of intense research on inbred strains of murine rodents, still there is no consensual answer to this question. In the context of adult neurogenesis, adulthood has been timed after weaning (at postnatal day 21), reasoning that by that age, the adult neurogenic niche, the subgranular zone (SGZ) of the dentate gyrus, has acquired its definitive structure (Kempermann, 2011); or more commonly, adulthood have been timed with sexual maturity, normally defined as achieving sexual reproduction and thus starting at puberty, around P30 (Kempermann, 2011). However, puberty is poorly correlated with successful mating, pregnancy, or maternal care, and it seems more reasonable to equate sexual maturity not just with the capacity for reproduction, but with the ability to reproduce and raise offspring successfully, that will be achieved at the end of the postpuberal adolescence period, a maturational stage that involves hormonal, physiological, neurological and behavioural changes that allow for successful reproduction (Arellano et al., 2024). Adolescence ends around P60 in both rats and mice, marking the beginning of adulthood (Arellano et al., 2024), and thus in this work we will consider P60 as the onset of adulthood for both species.

Since adulthood includes most of the lifespan in mammals, we will consider more meaningful subperiods: young adults, corresponding to peak reproduction (∼2-7 month-old for mice and ∼2-6 month-old for rats); middle age, corresponding to reproductive senescence in females (∼8-14 month-old in mice and ∼7-14 month-old in rats); and old age, corresponding to a post-reproductive (anestrous) period in females (∼15-30 month-old in mice and ∼15-21 month old in rats) as previously described (Arellano et al., 2024).

### The life history of new neurons in the adult dentate gyrus

To better understand the underpinnings of the model we propose, it is useful to review the general scheme of neurogenesis to define the specific phases of differentiation, functional integration, and maturation that new granule cells (GC) experience as they incorporate in the postnatal dentate gyrus. This is also important to define the terminology we will use along the manuscript to refer to new GCs at different developmental and maturational times. A scheme illustrating those process is shown in Fig 1.

**Figure 1.**
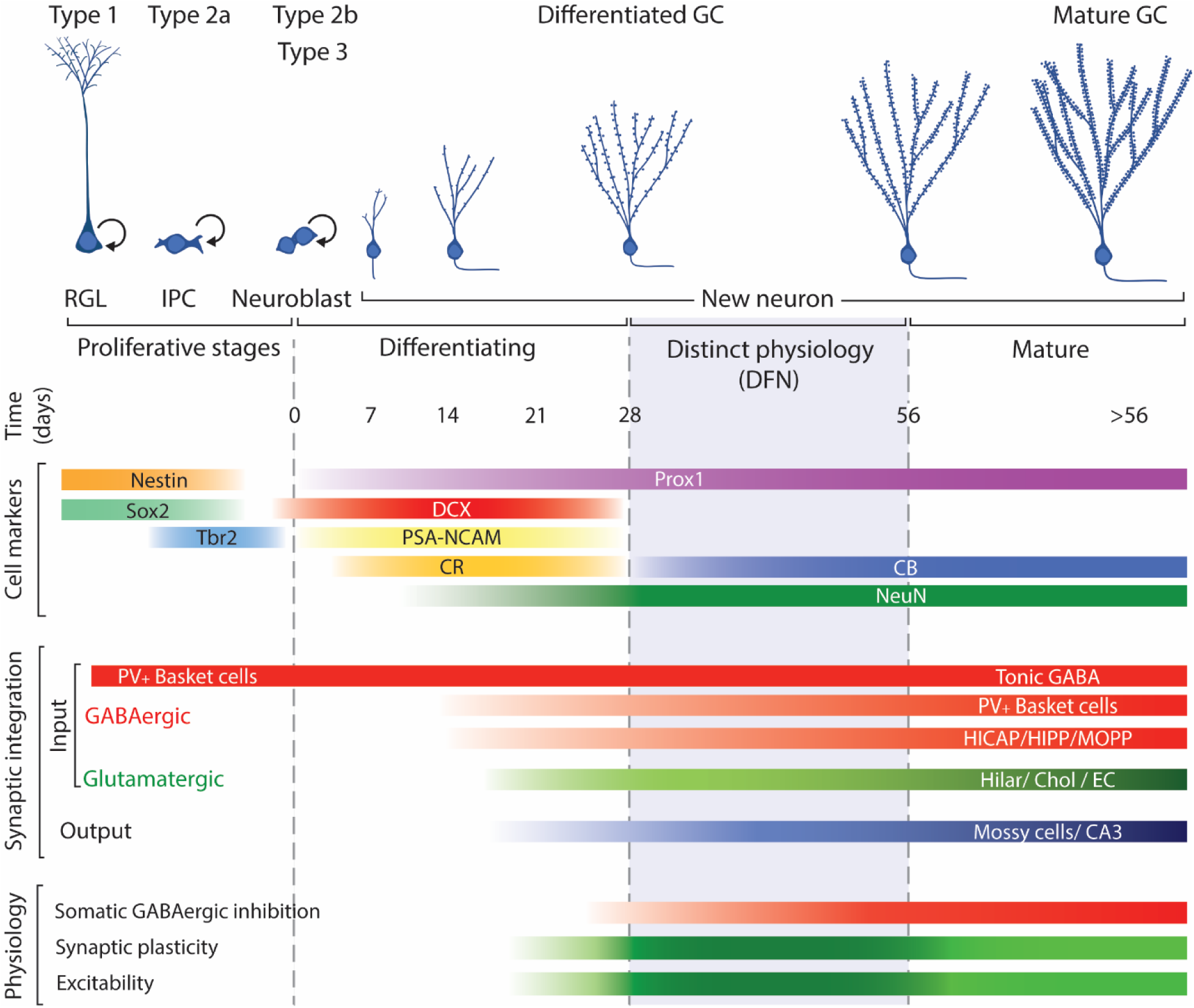
Schematic representation of the neurogenic process illustrating the main cell types involved: radial glia like progenitors (RGL; type 1 cells); intermediate progenitor cells (IPCs; Type 2a); neuroblasts, including committed neuronal IPCs (type 2b cells) and type 3 cells; and postmitotic neurons along their differentiation/maturation, timed in days. Cell markers include the approximate expression window along the neurogenic process of common markers used to identify each cell type. Synaptic integration summarizes the approximate timing when new neurons establish incoming GABAergic and glutamatergic input and output. Physiology reflects the neuronal properties of new neurons: darker green between 28 and 56 days reflects the enhanced plasticity and excitability of new neurons allowing them distinct functional capabilities while the subsequent lighter green represents normal mature physiological features. Modified from (Christian et al., 2014).

The accepted model of neurogenesis describes that type 1 radial glia-like progenitors in the subgranular zone of the dentate gyrus divide to produce type 2a cells, glia-like intermediate progenitors, that can divide and produce type 2b cells, neuronally committed intermediate progenitors that start expression of immature neuron markers DCX, polysialylated neural cell adhesion molecule (PSA-NCAM), and Calbindin-2, also known as Calretinin (CR; (Liu et al., 1996; Seki, 2002; Brown et al., 2003; Llorens-Martín et al., 2006; Plümpe et al., 2006; Spampanato et al., 2012). Type 2b progenitors can proliferate and produce type 3 cells that can include a small fraction of progenitors but is mostly composed of post-mitotic neurons that will develop to produce new GCs. Those post-mitotic neurons continue expression of DCX, PSA-NCAM, CR, and low levels of RBFox3, better known as neuronal nuclei (NeuN) that will increase as neuronal differentiation progresses (Seki, 2002; Brandt et al., 2003; Brown et al., 2003).

The differentiation of new neurons takes about 4 weeks, during which new neurons migrate to their final position, extend their dendrites and axons, establish synapses, and become functional (van Praag et al., 2002; Espósito et al., 2005; Zhao et al., 2006; Toni et al., 2007; Faulkner et al., 2008; Gu et al., 2012; Nakashiba et al., 2012; Vivar and van Praag, 2013; Jungenitz et al., 2014). Along that process of maturation, they down-regulate expression of immature neuron markers DCX, PSA-NCAM and CR while up-regulating expression of markers of mature GCs such as NeuN and Calbindin (CB; (Seki, 2002; Brandt et al., 2003; Brown et al., 2003; Rao and Shetty, 2004; Espósito et al., 2005; McDonald and Wojtowicz, 2005). During the differentiation phase, new neurons experience two main waves of cell death, the first at 2-3 days after cell division and the second 2-3 weeks into the differentiation process (Seki, 2002; Tashiro et al., 2006; Sierra et al., 2010). By 4 weeks of age, surviving new neurons have acquired GC morphology, are integrated in the hippocampal circuit and most of them have switched expression of DCX, PSA-NCAM and CR for NeuN and CB and survive for at least 6 months (Dayer et al., 2003; Kempermann et al., 2003; Rao et al., 2005; Morgenstern et al., 2008); although see (McDonald and Wojtowicz, 2005).

### Young new neurons do not have a different pattern of connectivity

Studies on young adult-born GCs have shown that they establish their connectivity following the pattern observed during dentate gyrus development (Espósito et al., 2005). At around two weeks of age they start receiving synaptic input from local GABAergic interneurons, although it produces depolarization, as young new neurons have high intracellular chloride concentration as their embryonic counterparts (Toni and Sultan, 2011). The next input are cholinergic afferents from septal areas, hypothalamus and basal forebrain, while the first glutamatergic synapses arrive during the third week from hilar mossy cells and entorhinal cortex and will grow rapidly through the following weeks. The last input to arrive is the GABAergic perisomatic inhibition, close to 4 weeks after mitosis (Espósito et al., 2005; Kumamoto et al., 2012; Vivar et al., 2012; Deshpande et al., 2013; Vivar and van Praag, 2013; Bergami et al., 2015). The output of new neurons also resembles that of mature granule cells, targeting hilar mossy cells, pyramidal neurons in CA3 and GABAergic interneurons in both the hilus and CA3. The axons of new GCs reach CA3 about 10-14 days after mitosis and start to exhibit sparse connections with CA3 neuron dendrites during the third week, although the mossy fiber extent, and the number of mossy terminals and their synapses continues growing for several weeks (Faulkner et al., 2008; Toni et al., 2008). Studies using viral labeling to identify single-synapse presynaptic targets of adult born neurons have shown atypical sources of input such as mature granule cells, CA3 or the subiculum. However, the input from mature granule cells has reasoned to be an artifact of the labeling technique (Deshpande et al., 2013; Bergami et al., 2015) while the projections from the subiculum and CA3 have been occasionally described in rats (Köhler, 1985; Li et al., 1994; Wittner et al., 2007), suggesting that they are not specific of adult born cells. Furthermore, those atypical sources of input to the DG are not consistently detected in all studies and represent a small fraction of the total labeled presynaptic. Therefore, it seems that adult born granule cells are not different in their pattern of connections compared to mature granule cells.

### Young new neurons 4 to 8 weeks old exhibit distinct physiology than mature GCs

Functional studies on new neurons have proposed they exhibit distinct physiological responses that confer them enhanced excitability and increased plasticity compared to mature GCs (Snyder et al., 2001; Schmidt-Hieber et al., 2004; Espósito et al., 2005; Ge et al., 2006, 2007; Overstreet-Wadiche et al., 2006; Piatti et al., 2006; Zhao et al., 2006; Marín-Burgin et al., 2012). Some studies have described physiological and functional critical windows in new neurons 1-3 weeks of age (Shors et al., 2001, 2002; Snyder et al., 2001, 2005; Jessberger and Kempermann, 2003; Madsen et al., 2003; Bruel-Jungerman et al., 2006; Tashiro et al., 2007; Deng et al., 2009; Trouche et al., 2009; Aasebø et al., 2011), although as described above, 1-3-week-old new neurons have not established enough afferent or efferent connections (van Praag et al., 2002; Zhao et al., 2006; Toni et al., 2007; Farioli-Vecchioli et al., 2008; Faulkner et al., 2008; Deshpande et al., 2013) to produce meaningful information processing in the hippocampal circuit. Furthermore, 14-day-old new neurons show dendritic extension reaching only to the IML (Perederiy et al., 2013), meaning they would not receive input from the EC terminating in the MML and OML, further suggesting they are not engaged functionally in a meaningful manner in the dentate circuit.

Alternatively, a more plausible window of enhanced plasticity and distinct functional properties has been described in new neurons between 4 and 8 weeks after cell division (Ge et al., 2007, 2008; Kee et al., 2007; Gu et al., 2012; Kheirbek et al., 2012; Kim et al., 2012; Marín-Burgin et al., 2012; Dieni et al., 2013; Brunner et al., 2014; Temprana et al., 2015; Danielson et al., 2016; McHugh et al., 2022; Mugnaini et al., 2023; Vyleta and Snyder, 2023), when new GCs are engaged in the hippocampal circuit (Espósito et al., 2005; Toni et al., 2007; Mongiat et al., 2009; Marín-Burgin et al., 2012). After that period, the physiology of new neurons becomes indistinguishable from that of mature GCs (Laplagne et al., 2006, 2007; Ge et al., 2007; Mongiat et al., 2009; Marín-Burgin et al., 2012; Brunner et al., 2014; Woods et al., 2018; McHugh et al., 2022; Mugnaini et al., 2023; Vyleta and Snyder, 2023), strongly suggesting that if new neurons have a distinct function of that of mature GCs, it must be performed during that time window, when they respond differently than mature cells. From now on, we will refer to new neurons during their first 4 weeks after mitosis as differentiating new neurons, while new neurons 4-8-week-old exhibiting the differential physiology conferring them enhanced plasticity and excitability will be referred as distinct functional neurons (DFNs). Adult-born neurons older than 8 weeks will be referred as mature new neurons or simply mature neurons, as they will be indistinguishable from the pool of GCs born during development of the animal (up to 2 months of age).

### How many new neurons are in the adult dentate gyrus?

Studies using 5-Bromo-2-deoxyuridine (BrdU) or Ki67 to assess proliferation have consistently revealed thousands of labeled cells in very young rats and mice (Cameron and McKay, 2001; Heine et al., 2004; McDonald and Wojtowicz, 2005; Snyder et al., 2009; Ben Abdallah et al., 2010), and it has been proposed that about half of those cells would survive and integrate in the dentate gyrus circuit, representing 5-6% of the rat GC population (West et al., 1991; Cameron and McKay, 2001; McDonald and Wojtowicz, 2005), prompting descriptions of thousands of new neurons produced every day in the rodent DG (e.g.: 4000-7000, (Kim et al., 2012); 9000, (Christian et al., 2014)). However, dentate neurogenesis in rodents decreases 15-20-fold by age 9-12 months (Kuhn et al., 1996; Bondolfi et al., 2004; Harrist et al., 2004; Heine et al., 2004; Hattiangady et al., 2008; Morgenstern et al., 2008; Ben Abdallah et al., 2010; Amrein et al., 2011; Encinas et al., 2011; Smith and Semënov, 2019), and therefore, the answer to the question of how many new neurons are available in the dentate gyrus will drastically depend on the age of the animal.

Studies evaluating the lifespan production of new neurons have shown more moderate figures. (Ninkovic et al., 2007; Imayoshi et al., 2008) used transgenic mice to label newly generated cells and reported values between 8-10% of the total population of 500,000 GCs (Kempermann et al., 1997a; Calhoun et al., 1998; Ben Abdallah et al., 2010; van Dijk et al., 2016), representing 40-50,000 new neurons added along adulthood. (Pilz et al., 2018) studied the process of neurogenesis in vivo and estimated that radial glial progenitors will produce on average 4.5 neurons that combined with the approximately 10,000 radial glial cells present in a 2-month-old mouse described by (Encinas et al., 2011) would mean that about 45,000 new neurons (or 9% of the total) are generated along the adult life of mice. (Lazic, 2012) used published data to estimate that C57 adult mice, defined as older than 3 months (Flurkey et al., 2007), would produce about 57,000 new neurons, representing 11.4% of the total number of GCs. (Smith and Semënov, 2019) proposed a model based on proliferation and cell cycle kinetics in C57 mice and predicted a total of 1.5 million new cells produced along the lifespan in both hemispheres (750,000 per hemisphere) with a 64-fold decline between 1-30 months of age. They do not specify how many of those would be new neurons, but if we assume about 40% (Kempermann et al., 1997b), then around 300,000 new neurons would be generated along the lifespan (per hemisphere), representing about 60% of the total granule cells. Such high levels of neurogenesis would imply the presence of neuronal replacement, as such large increase in the number of granule cells along the lifespan has not been reported.

From a functional point of view, however, we are not interested in the total number of new cells produced along life, but instead in the specific population of DFNs present at any given time in the dentate gyrus, as they might carry different functions than mature GCs. To the best of our knowledge, only Snyder and Cameron (Snyder and Cameron, 2012) have attempted to respond to this question before. They used a similar age criterion, (cells 8-weeks-old or younger) to define new neurons with a potential distinct functionality, although, as argued above, new neurons younger than 4 weeks might establish too few synapses to exert meaningful information processing in the hippocampal circuit (Espósito et al., 2005; Mongiat et al., 2009; Marín-Burgin et al., 2012). Nonetheless, Snyder and Cameron estimated that a 4-month-old rat would have about 170,000 young new neurons (per hemisphere) up to 8 weeks old, representing about 14% of the total number of GCs (Snyder and Cameron, 2012), while a 12-month-old rat would have about 25,000 new neurons 8-week-old or younger, representing 2% of the total. Overall, their model implied the generation of about 500,000 new neurons between ages 2-24 months, meaning about 40% of the population of GCs are replaced along the adult life of the animal (Snyder and Cameron, 2012). The Snyder lab updated the model recently predicting even higher levels of neurogenesis that represent about 50% of the total population of GCs along the lifespan of rats (Cole et al., 2020a). Overall, these models (Snyder and Cameron, 2012; Smith and Semënov, 2019; Cole et al., 2020b), predict much higher (∼5-fold) levels of neurogenesis than estimates based on empiric data (Ninkovic et al., 2007; Imayoshi et al., 2008; Pilz et al., 2018; Encinas et al., 2011; Lazic, 2012).

### A model to estimate the number of DFNs from the number of Doublecortin (DCX) labeled cells

We propose an alternative method to quantify the number of distinct functional neurons (DFNs) in the hippocampus based on the number of doublecortin (DCX) labeled cells. At face value, it seems reasonable to consider the number of cells expressing DCX as a measure of neurogenesis, since DCX is considered the canonical marker of newly generated neurons in the DG and has been proposed as a good estimator of the number of differentiating new neurons (Brown et al., 2003; Rao and Shetty, 2004; McDonald and Wojtowicz, 2005). However, it can be argued that although DCX is a good marker to obtain relative values of neurogenesis, for example to compare AHN across ages, experimental conditions or between species (Patzke et al., 2015), the population of DCX labeled cells might not provide an accurate estimate of the number of functional new neurons, as only a fraction of DCX labeled cells will survive to become fully functional (Plümpe et al., 2006). We can avoid the complexities of modelling those processes using empirical data to obtain the ratio between the number of differentiating new neurons and the number of DFNs 4-8 weeks old they produce. For this purpose, available data on BrdU injections with different survival times allows following a cohort of newly generated cells along their proliferation and differentiation process, to figure out the ratio between differentiating new neurons (type 3 cells labeled with BrdU and DCX/PSA-NCAM/CR) and the population of DFNs (labeled with BrdU and NeuN/CB). Once that ratio is known, the number of new differentiating neurons labeled with DCX present at any time will allow to estimate the population of DFNs.

Several reports have produced these data in rodents (Brandt et al., 2003; Brown et al., 2003; Kempermann et al., 2003; McDonald and Wojtowicz, 2005; Snyder et al., 2009), but we focused our attention on three of them (Brandt et al., 2003; Brown et al., 2003; McDonald and Wojtowicz, 2005), as the others included experimental manipulations (e.g.: multiple BrdU injections; use of non-naïve animals) that could affect the analysis (see suppl. data for details). From the selected studies, (Brown et al., 2003) only reported percentages of BrdU/DCX and BrdU/NeuN, so we combined their data with the number of cells BrdU positive reported by (McDonald and Wojtowicz, 2005) to use their dataset (**Table S1 and S2**). The graph in **Fig 2A** shows normalized data (percentage of the maximum) of BrdU/CR from (Brandt et al., 2003) in the mouse; BrdU/DCX in the rat from (Brown et al., 2003; McDonald and Wojtowicz, 2005) with their corresponding levels of BrdU/NeuN neurons 4-8 weeks. (Brandt et al., 2003) was performed in mice, but since all produced similar distributions of DCX and NeuN+ cells, we pooled their data with that of the rat studies.

**Figure 2.**
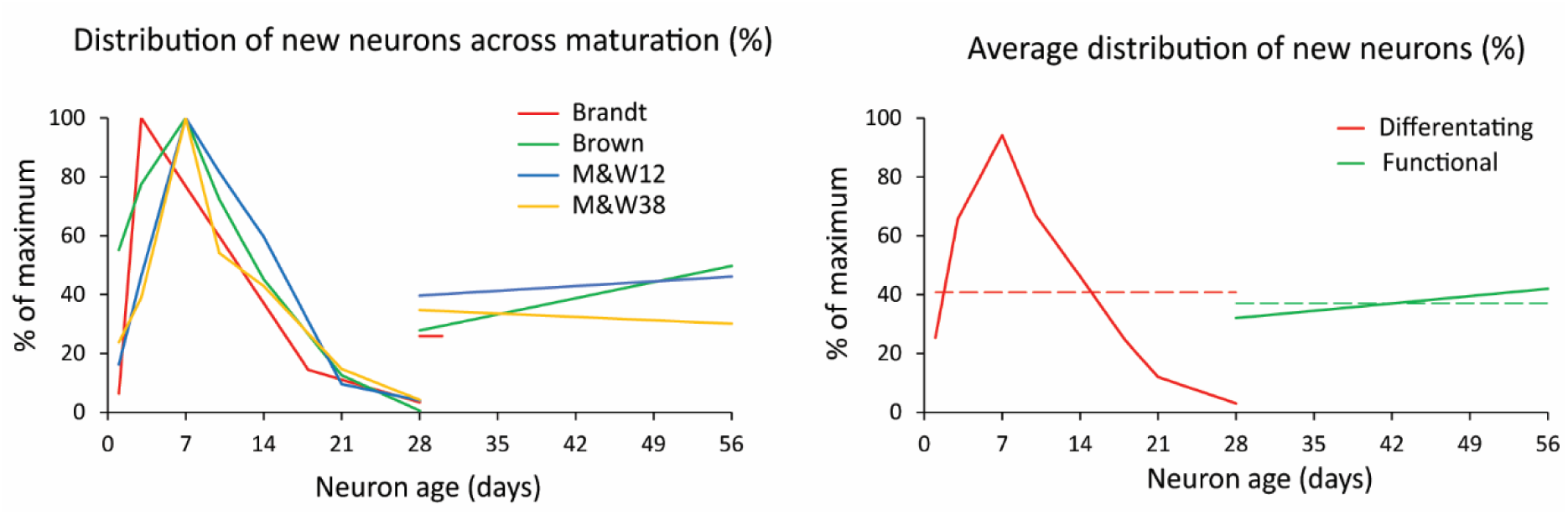
Distribution of new neurons across maturation. **A**. Normalized distribution of differentiating new cells (BrdU and DCX/CR labeling; % of the maximum) during the first 4 weeks after BrdU injection and normalized value of new functional cells (BrdU and NeuN/CB labeling) 4-8 weeks post injection. Differentiating cells exhibit a curve likely due to a combination of some delay in the onset of DCX/CR expression, proliferation and cell death that overall produce a considerable reduction of the initial population over 4 weeks. All data from rat except (Brandt et al., 2003) series in mouse. Note that Brandt et al., provided data on BrdU+/NeuN+ cells only at 28 days. **B.** Average of the normalized distributions shown in Fig 2A. Differentiating new neurons (BrdU and DCX/CR labeling) are represented in red and young functional neurons (BrdU and NeuN/CB labeling) in green. Dashed lines represent the average values of each population, 41% of maximum for differentiating neurons (red) and 37% for DFNs (green). The ratio between those values is 0.907 or 91% and represent the ratio between differentiating and functional new neurons from a cohort of newly generated cells (see suppl. data for extended explanation). M&W38 and 12 are abbreviations of P38 and 12-month-old rats in (McDonald and Wojtowicz, 2005).

The distributions showed that young neurons ranged between 26 and 43% of the maximum number of DCX+ cells, detected about 7 days post BrdU injection (**Table S1 and S2**). Given those similarities, we averaged the distributions to produce a single normalized dataset (**Fig 2B**) illustrating the distribution of differentiating new cells over time (red curve), and the distribution of functional new neurons 4-8 weeks old (green line). Those distributions summarize the populations of differentiating neurons and DFNs in a single cohort of new neurons. As both have the same duration (4 weeks), the ratio between the average value of those distributions (red and green dashed lines in Fig 2B) represents the ratio between the total population of differentiating cells and the total population of DFNs that will originate from them (for detailed explanation see suppl data). This ratio is 0.91, meaning that 91% of DCX labeled cells will become DFNs after an average period of 4 weeks.

Several reports have provided quantification of the total number of DCX labeled cells at different ages in rodents: (Rao et al., 2005, 2006; Zhang et al., 2008; Epp et al., 2009; Rennie et al., 2009) in rats and (Kronenberg et al., 2006; Ben Abdallah et al., 2010; Kuipers et al., 2015) 2015 in mice (**Fig 3A**). The distribution in rats shows more variability than in mice, with the datasets from Rao et al., and (Rennie et al., 2009) showing relatively high and low, respectively, numbers of DCX labeled cells, that combined might produce a balanced regression curve. However, (Rao et al., 2006) data decreased linearly with age, and even exhibits a slight increase in DCX labeled cells between 4 and 7 months of age, in contrast with the well-established notion of an exponential decline of neurogenic parameters across age (Kuhn et al., 1996; Heine et al., 2004; Amrein et al., 2011) and therefore, the data from (Rao et al., 2006) were not used. The number of DCX labeled cells in rat are approximately double than those reported in mice, as expected since rats have double number of GCs than mice (Boss et al., 1985; West et al., 1991; Hosseini-Sharifabad and Nyengaard, 2007; Fitting et al., 2010).

**Figure 3.**
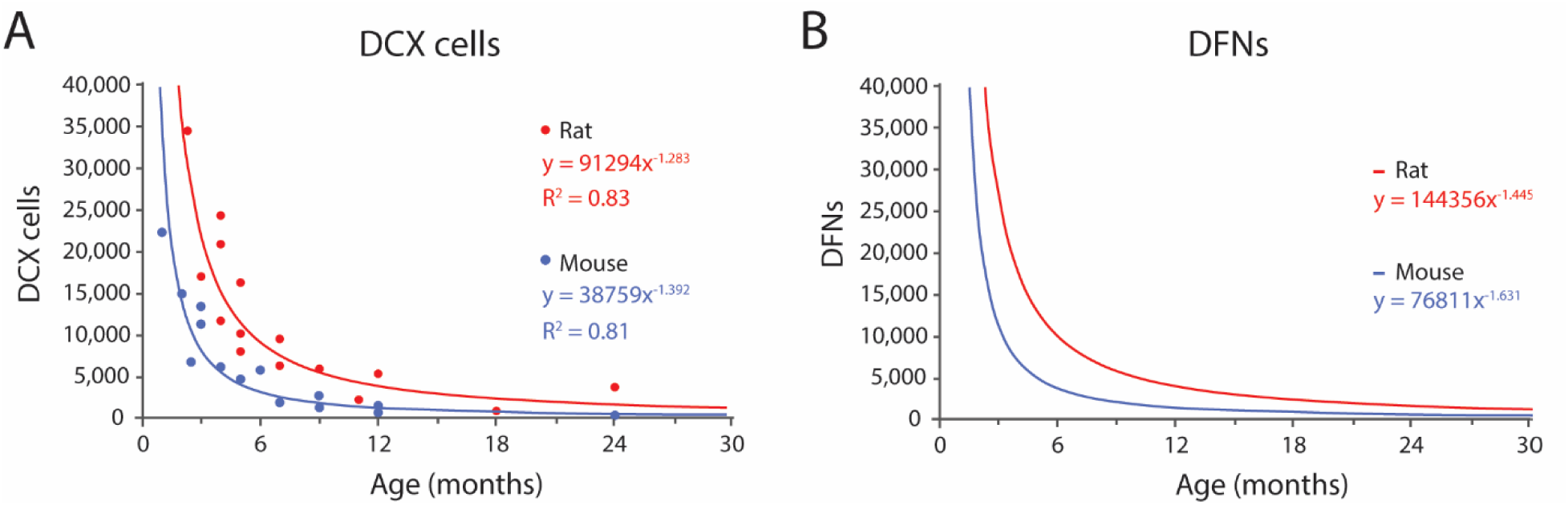
Modeled distributions of DCX labeled cells and DFNs in mice and rats across ages. **A,** Distribution of DCX labeled cells in rats and mice across age according to published results (see text for references). Both distributions fit negative power equations. **B**, Distribution of DFNs in rats and mice across age from regression of DCX datasets transformed by multiplying each value by 0.91 (survival rate) and adding one month to the age to account for differentiation delay. The resulting values fit negative potential distributions. See Table 1 for values at selected ages.

In mice, (Kronenberg et al., 2006; Ben Abdallah et al., 2010; Kuipers et al., 2015) provided different time points between 1 to 24 months of age. Their combined data fit best an exponential distribution (R^2^=0.84), that however models very poorly the data for young ages, and we selected instead a power distribution that had slightly lower fit (R^2^=0.81) but much better predictive value at young ages, when the levels of neurogenesis are the highest (for details see suppl. data). The resulting equation for mice is:

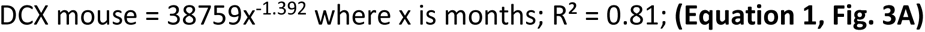

In rats, (Rao et al., 2005; Zhang et al., 2008; Epp et al., 2009; Rennie et al., 2009) provided timepoints between 2.3 and 24 months. In this case, the data fitted best a power distribution with equation:

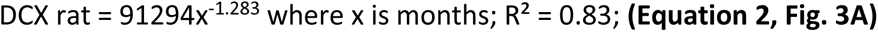

Both equations describe the number of DCX labeled cells along time, and 91% of those will become DFNs after an average 4-week differentiation period. Thus, the number of DFNs at a given time, correspond to 91% (survival rate) of the number of DCX labeled cells 4 weeks before (see suppl data for detailed explanation). The estimated values of DFNs can be modeled with power distributions as follows:

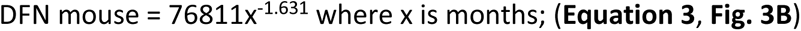

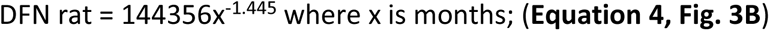

The modeled number of DCX labeled cells, DFNs and their percentages from the total population of GCs are shown in **Table 1A and 1B** for mice and rats respectively. Total population was estimated to be 500,000 GCs in the mouse (Kempermann et al., 1997a; Calhoun et al., 1998; Ben Abdallah et al., 2010; van Dijk et al., 2016) and about 1,000,000 GCs in the rat (Boss et al., 1985; West et al., 1991; Hosseini-Sharifabad and Nyengaard, 2007; Fitting et al., 2010); data per hemisphere).

**Table 1A.** Main parameters of neurogenesis in mice along age according to the model. See legend under Table 1B.

| Mouse |  | DCX+ cells |  | DFNs |  |  |
| --- | --- | --- | --- | --- | --- | --- |
| Life stage | Age (months) | N | % of total GCs | N | % of total GCs | % activated GCs |
| developmental | 1 | 38,759 | 7.8 | - | - | - |
|  | 2 | 14,769 | 3.0 | 35,271 | 7.1 | 29.7 |
| Young adult | 3 | 8,399 | 1.7 | 13,439 | 2.7 | 13.9 |
|  | 4 | 5,627 | 1.1 | 7,643 | 1.5 | 8.4 |
|  | 6 | 3,200 | 0.6 | 3,754 | 0.8 | 4.3 |
|  | 7 | 2,582 | 0.5 | 2,912 | 0.6 | 3.4 |
| Middle age | 8 | 2,144 | 0.4 | 2,350 | 0.5 | 2.7 |
|  | 12 | 1,219 | 0.2 | 1,253 | 0.3 | 1.5 |
|  | 14 | 984 | 0.2 | 993 | 0.2 | 1.2 |
| Old age | 15 | 894 | 0.2 | 895 | 0.2 | 1.1 |
|  | 24 | 465 | 0.1 | 449 | 0.1 | 0.5 |
|  | 30 | 341 | 0.1 | 325 | 0.1 | 0.4 |

**Table 1B.**
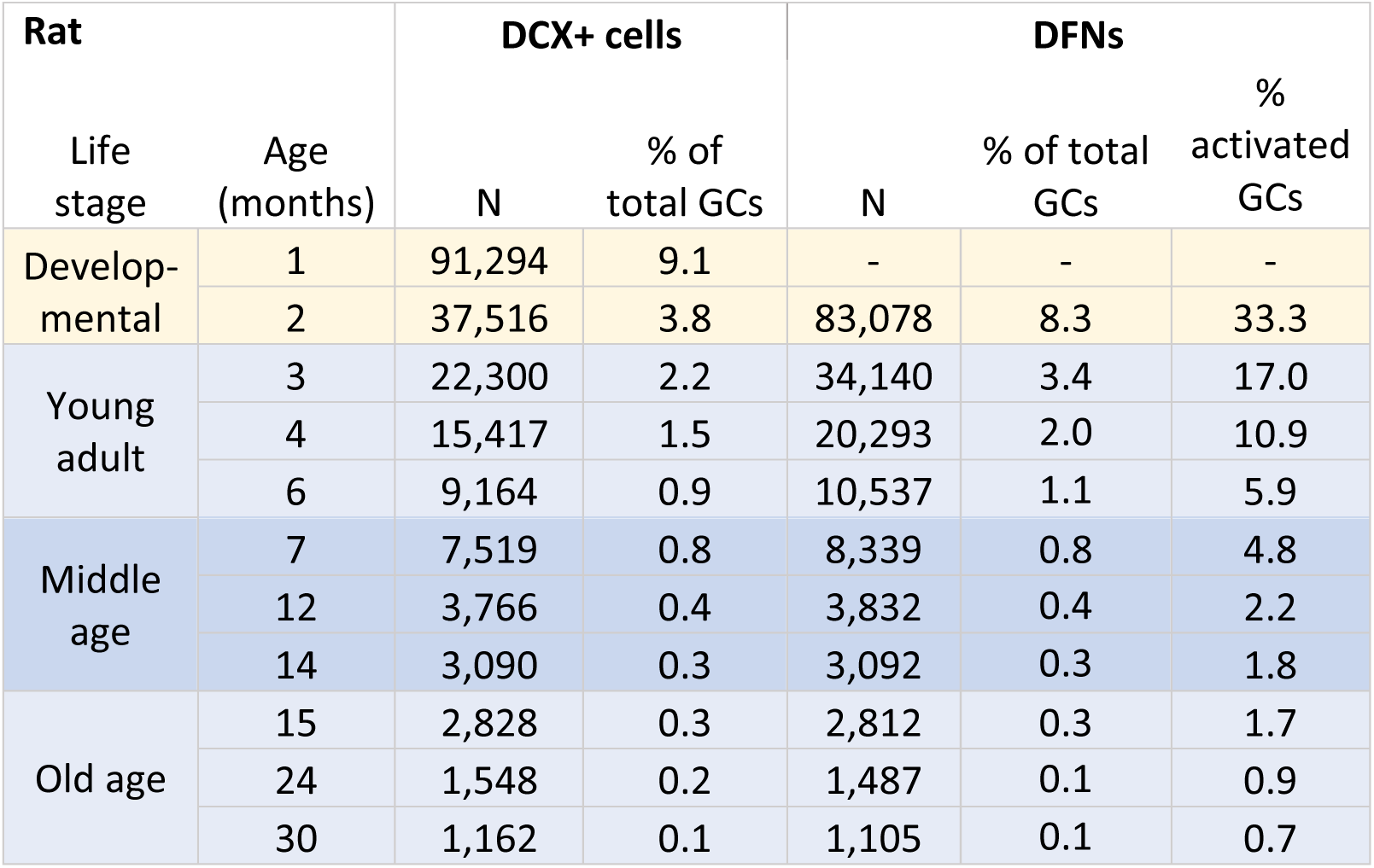
Main parameters of neurogenesis in rats along age according to the model. Yellow rows represent developmental neurogenesis, while blue cells represent adult neurogenesis (neurons born in 2-month-old or older animals) in young, middle age and old animals (Arellano et al., 2024). DCX values are derived from equations 1 and 2 for mice and rats respectively (see text). The number of DFNs at any given time is 0.91 of the DCX labeled cells present 4 weeks before. Note the precipitous decline in neurogenesis in the first months. Percentage of new neurons calculated over the total population of GCs (∼500,000 in mouse and ∼1,000,000 in rats). Percentage of activated GCs describes the fraction of GCs activated by stimulation of the perforant pathway that are DFNs (from data collected in mice by (Marín-Burgin et al., 2012).

| Rat |  | DCX+ cells |  | DFNs |  |  |
| --- | --- | --- | --- | --- | --- | --- |
| Life stage | Age (months) | N | % of total GCs | N | % of total GCs | % activated GCs |
| Developmental | 1 | 91,294 | 9.1 | - | - | - |
|  | 2 | 37,516 | 3.8 | 83,078 | 8.3 | 33.3 |
| Young adult | 3 | 22,300 | 2.2 | 34,140 | 3.4 | 17.0 |
|  | 4 | 15,417 | 1.5 | 20,293 | 2.0 | 10.9 |
|  | 6 | 9,164 | 0.9 | 10,537 | 1.1 | 5.9 |
| Middle age | 7 | 7,519 | 0.8 | 8,339 | 0.8 | 4.8 |
|  | 12 | 3,766 | 0.4 | 3,832 | 0.4 | 2.2 |
|  | 14 | 3,090 | 0.3 | 3,092 | 0.3 | 1.8 |
| Old age | 15 | 2,828 | 0.3 | 2,812 | 0.3 | 1.7 |
|  | 24 | 1,548 | 0.2 | 1,487 | 0.1 | 0.9 |
|  | 30 | 1,162 | 0.1 | 1,105 | 0.1 | 0.7 |

It is important to note that adult neurogenesis, by definition, implies neurons born in adult animals. Since adulthood is achieved around 2 months of age, new neurons born at that age will only become functional when the animal is 3 months old, after the 4-week differentiation period. In turn, the cohort of DFNs present when the animal turns adult, at P60, was generated around P30, and correspond to adolescent neurogenesis (yellow cells in **Table 1**). Thus, we will refer to adult born, DFNs as those present in the dentate gyrus at 3 months or later (blue cells in **Table 1**). Both mice and rats show similar proportion of DFNs across age. In mice, the model shows a peak population of 13,500 DFNs (2.7% of the total population) in 3-month-old animals, that declines very rapidly: by the beginning of middle age (∼8 months) there are about 2,500 DFNs (0.5%), and by the onset of old age (∼15 months) there are about 900 DFNs (0.18%; **Table 1A**). Rats show similar distributions, and at 3 months the model predicts about 34,000 DFNs (3.4% of the total) that decreases rapidly to about 7,500 (0.8 %) in early middle age (∼7 months) and about 2,800 (0.28%) at the onset of old age (∼15 months; **Table 1**). Comparison between the distribution of DFNs in mice and rats reveal very similar curves, both fitting well power distributions, obviously with higher values for rats, given they have double the number of GCs in the dentate gyrus. Regarding the proportion of DFNs over the total population, young rats exhibit a higher proportion (30% more) than mice, a difference that grows with age to reach 60% more in old rats (15 months and older) compared to mice.

Integration of equations 3 and 4 allows estimating the number of new neurons produced at any interval of time in both species, showing that during the adult life of a C57 mouse (2-30 months of age; Arellano et al., 2024) about 47,000 new neurons will be produced, ∼10% of the total GC population. In rats, about 115,000 new neurons or ∼12% of the total population are produced between age 2-21 months. Note that the integration should be done between 3-30 months, as there are no adult born DFNs at 2 months of age. As expected, the model predicts that most of those new neurons, 76% in mice and 79% in rats will be produced in the first year, a clear reminder of the exponential decline of neurogenesis with age.

### Preferential recruitment of new neurons

It has been argued that the low number of DFNs can be counterbalanced by their preferential activation in the dentate network, at a 2-6-fold rate compared with mature GCs, so sufficient new neurons may be recruited to have a relevant functional contribution (Ramírez-Amaya et al., 2005; Kee et al., 2007; Marín-Burgin et al., 2012). Specifically, (Marín-Burgin et al., 2012) reported an example of a single stimulus that recruited 5% mature neurons but 30% DFNs in mice about 3-month-old, suggesting a prominent role for new neurons to activate CA3. When those ratios are transformed into absolute numbers, excitation of the DG will activate ∼25,000 mature neurons (5% of the total 500,000 GCs in mice), and ∼4,000 DFNs (30% of 13,500 DFNs) indicating that DFNs represent about 14% of the activated GCs in a 3-month-old animal (**Table 1A**). If this proportion holds across age, given the exponential decrease of DFNs, less than 3% DFNs will be activated in a middle-aged mouse and ∼1% in an old animal (**Table 1A**). An alternative possibility is that the number of activated DFNs described by (Marín-Burgin et al., 2012) is maintained across age, meaning around 4000 DFNs can be activated by a single stimulus at all ages. However, the number of DFNs decreases below 4,000 around 6 months of age (**Table 1A**) meaning that in older animals a much lower proportion of DFNs will be activated by a single stimulus. Extrapolation of those data to rats shows similar percentages of activated DFNs in young adults and about double ∼5% and ∼2% in middle age and old animals respectively (**Table 1B**), still representing low proportions of the total population of activated granule cells.

## Discussion

The high levels of neurogenesis in the young rodent hippocampus have been central to discussions on its potential function for the last 3 decades. The hippocampus is involved in a large variety of functions and numerous reports have proposed a functional involvement of adult neurogenesis in virtually all those functions (Kim et al., 2012; Snyder and Drew, 2020; Kempermann, 2022). From a computational point of view, three main conditions need to be satisfied for new neurons to have a functional impact. First, new neurons must exhibit differential connectivity and/or functional responses from mature cells; second, there needs to be enough new neurons to have a measurable contribution, and third, those new neurons need to have enough input and output in the hippocampal network to carry out their function. In this meta-analysis we attempted to answer the first two questions by creating a model to estimate the number of new GCs with differential functional capabilities across age in mice and rats. Due to space constrains, we do not address the third question that will be analyzed in a separate manuscript.

### The differential physiology of adult generated GCs

Available data indicates that while new granule cells exhibit similar pattern of connectivity to that of mature granule cells born during perinatal development, adult born neurons show differential excitability and plasticity compared to mature GCs. Two main periods have been described: between 1 and 3 weeks of age, and between 4-8 weeks of age.

The first period was described by studies using ablation of new neurons through X-ray irradiation or pharmacological interventions, showing that new neurons seemed to be required for different hippocampal functions between 1-3 weeks after mitosis (Shors et al., 2001, 2002; Snyder et al., 2001, 2005; Jessberger and Kempermann, 2003; Madsen et al., 2003; Bruel-Jungerman et al., 2006; Tashiro et al., 2007; Deng et al., 2009; Trouche et al., 2009; Aasebø et al., 2011). However, at those ages new neurons exhibit short dendritic arbors and sparse synaptic input mostly from local GABAergic interneurons that is however excitatory in nature (Espósito et al., 2005; Toni and Sultan, 2011), and start to receive input from hilar mossy cells and entorhinal cortex (Espósito et al., 2005; Kumamoto et al., 2012; Vivar et al., 2012; Deshpande et al., 2013; Vivar and van Praag, 2013; Bergami et al., 2015). Similarly, the axons of new GCs reach CA3 about 10-14 days after mitosis, but only sparse, immature synapses are found at 3 weeks after mitosis (Faulkner et al., 2008; Toni et al., 2008). Thus, new neurons at those early ages are not fully engaged in the hippocampal network, as they barely receive entorhinal input or convey excitatory output to CA3. Furthermore, the excitatory effect of their GABAergic input implies they lack inhibition and therefore their excitability might not be modulated by the hippocampal network, suggesting they might be a source of noise more than a computational asset in the dentate gyrus network.

A second body of evidence that has become the mainstream view on the field has showed that new neurons 4-8-weeks-old exhibit enhanced physiological features compatible with increased excitability and plasticity (Ge et al., 2007, 2008; Kee et al., 2007; Gu et al., 2012; Kheirbek et al., 2012; Kim et al., 2012; Marín-Burgin et al., 2012; Dieni et al., 2013; Brunner et al., 2014; Temprana et al., 2015; Danielson et al., 2016; McHugh et al., 2022; Mugnaini et al., 2023; Vyleta and Snyder, 2023). This window seems more realistic, since there is consensus that 4-week-old new neurons are integrated in the hippocampal circuit, able to respond to different input sources including the entorhinal cortex and capable of producing synaptic output to mossy cells and CA3 pyramidal neurons (van Praag et al., 2002; Espósito et al., 2005; Zhao et al., 2006; Toni et al., 2007; Faulkner et al., 2008; Gu et al., 2012; Nakashiba et al., 2012; Vivar and van Praag, 2013; Jungenitz et al., 2014; Lopez-Rojas and Kreutz, 2016; Mugnaini et al., 2023). Based on these data, we considered 4-8-weeks-old new neurons as distinct functional neurons or DFNs, the relevant population to quantify to assess the potential for a functional role of AHN. As described earlier, when we refer to mature GCs we include both developmentally generated (born in animals up to 2 months of age) and adult-born neurons older than 8 weeks, that share similar physiological responses.

Although the 4-8 weeks critical window has emerged as the most widely accepted in the field, different studies provide different timing. For example, the most widely cited of those physiological studies, (Ge et al., 2007) and others (Espósito et al., 2005; Mongiat et al., 2009; Restivo et al., 2015; Temprana et al., 2015) including behavioral studies (Denny et al., 2012) describe a critical period restricted to 4-6-week-old neurons, with 7-weeks-old and older new neurons responding similar to mature GCs. In this scenario, the proportion of DFNs reflected in **Table 1** would be reduced by half at any age and would mean that a 3-month-old mouse would have about 1.3% (∼7,000) DFNs, while middle age and old mice would have about 0.25% (∼1,200) and ∼0.1% (about 450) DFNs, very little fractions of new neurons whose consequences will be discussed in the next sections.

### The model predicts a low number of DFNs in the adult dentate gyrus, particularly in middle aged and old animals

Our model relates the number of DFNs to the population of DCX labeled cells in the dentate gyrus, allowing to use published data on the distribution of DCX labeled cells across age to estimate the number of DFNs. The model predicts that about 91% of the DCX labeled present at one point will become DFNs 4 weeks later. We pooled data from studies using stereology to report the absolute number of DCX labeled cells across age both in mouse (Kronenberg et al., 2006; Ben Abdallah et al., 2010; Kuipers et al., 2015) and in rat (Rao et al., 2005; Zhang et al., 2008; Epp et al., 2009; Rennie et al., 2009) to describe the distribution of DFNs across age in both mice and rats. An advantage of relying on DCX to estimate the number of new neurons is that immunolabeling is a priori more consistent between experiments and labs than BrdU injections, which vary widely depending on dosage, timing of post-injection analysis, differences in clearance of the marker and potential dilution of BrdU labeling (Smith and Semënov, 2019). In support of this point, the pooled data of DCX labeled cells from different studies in both species produce coherent datasets fitting negative power distributions with high regression coefficients. Although the distribution of DCX labeled cells in mice fitted best an exponential distribution (R^2^=0.84), the resulting curve modeled very poorly the data for young ages (**Table S3**), and we selected instead a power distribution that had slightly lower correlation coefficient (R^2^=0.81) but much better predictive value at young ages, when neurogenesis is the highest (**Table S3**). The negative power functions modeling DCX labeled cells can be hypothesized to have their maximum at the peak of developmental neurogenesis around P7 (Angevine, 1965; Schlessinger et al., 1975; Bayer, 1980).

The distribution of DFNs across age were obtained after transforming the DCX data according to the parameters obtained from the model: 0.91% survival rate and 4-week differentiation delay, resulting also in negative power distributions (**Fig 3B**). We set the onset of adulthood at 2 months of age (Arellano et al., 2024), meaning that DFNs will only be available at 3 months of age, as new neurons born in 2-month-old animals will take about 4 weeks to differentiate and become functional. The 91% survival rate suggest that both populations (DCX labeled cells and DFNs) are similar in size and would seem to confirm the generally accepted idea that DCX is a good reflection of the number of new neurons in the dentate gyrus or mice and rats (Brown et al., 2003; Rao and Shetty, 2004; McDonald and Wojtowicz, 2005; Ben Abdallah et al., 2010; Patzke et al., 2015; Lipp, 2017). Comparison of both populations show similar values in middle age and old individuals, but significant differences in very young animals (1.6-fold, **Table 1**) due to the sharp decline of the DCX population at early ages. Therefore, the assumption that DCX is a good reflection of the number of new neurons in the dentate gyrus is valid for middle-aged and old animals, but only allows a rough estimate of the number of DFNs in young adults, that will have more DFNs than DCX labeled cells. In this regard, it can be argued that we have not considered possible differences in fate choice, pace of maturation or survival of new neurons with age. However, most studies have pointed no differences in such parameters (McDonald and Wojtowicz, 2005; Rao et al., 2005, 2006; Hattiangady and Shetty, 2008) and the datasets in our model comparing very young (adolescent) P38 with aged 12-month-old animals only showed differences in the rate of new neuron survival, that was slightly lower in older animals.

The model shows similar distribution of DFNs in mice and rats, both fitting power curves, obviously with higher values for rats as they have double the number of GCs in the dentate gyrus. The proportion of DFNs over the total population in both species show very similar ratios in young animals that start to diverge as animals age, with rats exhibiting larger proportion of DFNs compared to mice, reaching a 1.6-fold difference after 12 months of age. According to the model, 3-month-old animals have about 3% DFNs in both species, quite a low percentage considering this will be the peak of the DFN population. As age increases, the number of DFNs decreases drastically, and by the onset of middle age (7-8 months), both species show less than 1% (0.5-0.8%) DFNs over the total population of GCs, that decreases to very low levels (0.2-0.3%) by the beginning of old age at ∼15 months (Table 1).

### Our model in context: agreement with experimental evidence; disagreement with proliferation-based models

The model described here predicts that along the adult lifetime (2-30 months of age) of C57Bl/6 mice, the most common strain used in research, about 47,000 new neurons are produced, that represent ∼10% of the total population of GCs. Regarding rats, the model predicts that along adulthood (2-21 months) rats produce about 115,000 new neurons, or ∼12% of the total GCs, a slightly higher proportion than in mice, as previously suggested (Snyder et al., 2009). The value for mice is in good agreement with previously reported values of 8-11% using gene reporters (Ninkovic et al., 2007; Imayoshi et al., 2008), combining data on neural progenitors and their expected neuronal offspring (Pilz et al., 2018; Encinas et al., 2011) or from models based on empiric data on neurogenesis markers (Lazic, 2012).

Conversely, models based on proliferation data and relying on estimates of cell cycle duration and frequency of progenitor division have predicted much higher proportion of new neurons along the lifespan, from 40-50% of the total GC population in rats (Snyder and Cameron, 2012; Cole et al., 2020a) to ∼60% in mice (Smith and Semënov, 2019). These high estimates are in clear disagreement with our results and with previous experimental evidence reported above (Ninkovic et al., 2007; Imayoshi et al., 2008; Encinas et al., 2011; Pilz et al., 2018), and a pivotal question is how those studies reconcile such large number of newly generated neurons with much low number of DCX labeled cells present in mice (Kronenberg et al., 2006; Ben Abdallah et al., 2010; Kuipers et al., 2015) or rats (Rao et al., 2005; Zhang et al., 2008; Epp et al., 2009; Rennie et al., 2009). For example, the model described by (Cole et al., 2020a) predicts ∼175,000 new neurons generated between 2 and 3 months of age in rats (equation in Fig. 8A in (Cole et al., 2020a) although see note in Suppl. data). Thus, one would expect similar or slightly higher number of DCX labeled cells (∼175,000) at around 2 months of age, but instead, a 2.3-month-old rat has less than 35,000 DCX labeled cells (Epp et al., 2009), about five times less than expected from the model. Therefore, those high levels of neurogenesis conflict with the available evidence and more generally, with the levels of neurogenesis suggested by DCX labeled cells in rodents. And, at those elevated rates of neurogenesis, neuronal replacement would be so high that the risk of disruption and interference in memory processes might surpass the benefit of adding new, more responsive neurons to the network (Rakic, 1985; Akers et al., 2014).

Our mice model is based mostly on data from C57 animals, and (Kempermann et al., 1997b) showed that 2-month-old C57 mice injected with BrdU for 6 days showed about 0.36% labeled GCs 4 weeks later, suggesting that at that rate, about 1.8% of GCs would be generated between 2 and 3 months. Our results in mice point to higher levels of neurogenesis (2.7%, **Table 1A**), indicating the model does not seem to be biased to minimize AHN. Additionally, it is important to contextualize the results of our model into the inter-strain variability documented in mice and rats. (Kempermann et al., 1997b) also reported that 129/SvJ exhibited about half new neurons after 4 weeks than C57, BALB/c or CD1 mice. This implies that the predicted numbers of DFNs revealed by our model would be halved for 129/SvJ mice. Considering the proposed essential role of AHN in hippocampal function, this remarkable inter-strain difference suggests 129/SvJ mice might exhibit some cognitive phenotype regarding hippocampal function. However, as far as we know, there are no reports of this strain as less capable cognitively, showing accelerated cognitive decline with age or predisposition to hippocampal function impairment that would be expected considering the extremely low levels of DFNs predicted for that strain in middle-aged and old animals (∼0.25% of the total GCs at middle age).

### The limitations imposed by DFNs low numbers and the role of age and preferential activation

The proportions of DFNs in the adult dentate gyrus reported here represent, objectively, a very small fraction of the total population of granule cells even in very young individuals. Such lower levels of DFNs challenge the notion that new neurons have an essential contribution in the dentate gyrus circuit and are required for proper hippocampal function, a concept that has been central in the field (Snyder and Drew, 2020; Kempermann, 2022). Shifting the perspective to mature granule cells, is hard to explain how, when adult neurogenesis is compromised, the remaining ∼97% of mature granule cells are incapable of performing their normal function in the hippocampus. This conundrum escalates when we consider that the dentate gyrus of middle age and old animals contains more than 99% mature granule cells, still unable to deliver normal dentate gyrus function. And as indicated when discussing the critical period of new neurons, if the critical window is defined as 4-6 weeks of age as proposed by several studies (Ge et al., 2007; Espósito et al., 2005; Mongiat et al., 2009; Restivo et al., 2015; Temprana et al., 2015; Denny et al., 2012) then the fraction of DFNs is reduced by half, deepening the physical and logical barrier for an essential role of new neurons.

The functional limitations imposed by those low numbers have been challenged by studies describing preferential activation of DFNs due to their enhanced excitability during normal activity of the dentate gyrus (Ramírez-Amaya et al., 2005; Kee et al., 2007; Marín-Burgin et al., 2012), although there is not clear consensus in this point, as other studies could not identify such preferential activation, showing similar activation of DFNs and mature GCs (Stone et al., 2011; Dieni et al., 2013). Our analysis shows that when preferential recruitment of DFNs is translated to actual numbers using Marin-Burgin et al., data, the maximum contribution of DFNs to the pool of activated GCs would be 14% in 3-month-old mice. This proportion is relatively high and seems compatible with a more relevant functional contribution of DFNs in the activity of the network, but still indicates that the vast majority (86%) of the output to mossy cells, CA3 and GABAergic interneurons will be driven by mature GCs (**Table 1**). Under the same paradigm, during normal dentate gyrus activation in middle age and old animals, only ∼3% and ∼1% of activated granule cells will be DFNs, respectively (**Table 1**), meaning that at those ages over 97% of cells responding to a given stimulus in the dentate gyrus would be mature GCs, arguing against the idea that mature GCs “retire” to let new neurons take care of business in the dentate gyrus (Alme et al., 2010), and more importantly, implying that even with preferential activation, the number of activated DFNs in older animals is very low. Furthermore, the drop in the number DFNs with age is so pronounced that even if all DFNs present in the DG of middle-aged and old animals are activated by a given stimulus, they would represent only ∼10% and ∼4% of the total activated cells at those ages. In summary, even with preferential recruitment, the number of activated DFNs represents a small fraction of activated granule cells, once again questioning if such low levels of activation can support the essential role in hippocampal function consistently reported, particularly in middle aged and old animals.

As previously stressed (Snyder, 2019; Arellano et al., 2024), many of the functional studies supporting an important role for new neurons used very young, in many cases adolescent animals, that exhibit larger rates of neurogenesis, up to 7-8% according to our model (**Table 1**). If we consider the potential contribution of preferential recruitment of DFNs at those early ages, the model indicates that up to 30% of DFNs could be activated during normal DG activity (although see (Stone et al., 2011; Dieni et al., 2013)). In those conditions, it is reasonable to consider that DFNs might have a relevant, maybe critical participation in the hippocampal network, that however will quickly diminish as the levels of neurogenesis decrease rapidly in the following months. This potential bias introduced by studies in very young animals might partially explain the apparent conundrum between the low number of DFNs and their proposed essential function. A reasonable strategy to solve this issue would be to perform studies in middle-aged animals, that should provide a more balanced assessment, not only to understand the possible role of neurogenesis in rodents, but also to establish more realistic and useful comparisons with humans.

### Long term functional differences of adult-born GCs

Given the potential functional limitations of DFNs imposed by their low numbers in aging animals, it has been proposed that adult born GCs exhibit different properties than mature, developmentally generated GCs, beyond the 4-8 weeks of age when they exhibit enhanced physiology and plasticity (Lemaire et al., 2012; Snyder, 2019; Cole et al., 2020a). Those claims are based on results in rats showing extended structural plasticity in the form of dendritic remodeling in response to activity (Lemaire et al., 2012) and increase in input connectivity reflected by higher numbers of dendritic spines (Cole et al., 2020a). At face value, increased connectivity of adult born neurons might facilitate their recruitment and information processing in the dentate network. However, the studies on the physiology of adult born neurons coincide in describing new neurons beyond 7-8 weeks old as physiologically indistinguishable from mature GCs (Laplagne et al., 2006, 2007; Ge et al., 2007; Mongiat et al., 2009; Marín-Burgin et al., 2012; Brunner et al., 2014; Woods et al., 2018; McHugh et al., 2022; Mugnaini et al., 2023) and therefore, is hard to justify how their activity would be different from that of mature GCs. Another implication of a long-term effect of adult born neurons is that their functional contribution should increase over time as their number grows, a notion difficult to reconcile with the well-known decline in cognitive function related to aging.

## Conclusion

There seems to be an overall assumption in the field of AHN, that new, adult-born neurons and their increased excitability and plasticity are essential for the normal function of the hippocampus. AHN has been related to most if not all hippocampal functions such as memory consolidation, reward learning, emotional contextualization, time stamping or even forgetting (Kempermann, 2022) and alterations in AHN have been related to cognitive deficits associated to old age, neurodegenerative diseases such as Alzheimer’s and in many other neuropsychiatric disorders, from depression to autism (Choi and Tanzi, 2019; Kerloch et al., 2019; Babcock et al., 2021; Bicker et al., 2021; Sun et al., 2022; Tartt et al., 2022).

The results of our analysis showing low numbers of functionally distinct new neurons in the dentate network and their small rates of activation during normal stimulation, challenges the notion of their prominent functional role in the adult rodent hippocampus, particularly in middle age and old animals. This controversy might be in part explained because most functional studies might be biased since they have been performed in very young, sometimes adolescent, rats and mice when they exhibit peak neurogenesis, disregarding the much lower levels of new neurons present in middle aged or old rodents (Snyder, 2019; Arellano et al., 2024). For the same reason, extrapolation of those data to humans might not be very useful, as there might not be much interest in improving cognition of people in their 10s and 20s when they are in their cognitive prime, while it could be relevant to help people say beyond their 60s and 70s, when hippocampal function might take a hit due to aging or neurological disease.

As described in the introduction, three main questions must be answered to evaluate the potential functional role of new neurons. We have addressed the first two questions in this manuscript, revealing low numbers of new neurons with differential physiology. The remaining question is if those new neurons have enough connectivity during their critical period to support such differential function. Several studies have indicated that they are fully integrated in the hippocampal circuit, but their connectivity is still developing and differs from that of mature granule cells in terms of number of synaptic contacts and maturity of those connections (van Praag et al., 2002; Toni et al., 2007, 2008; Faulkner et al., 2008). Detailed analysis of this question cannot be included in this manuscript due to space constraints and will be analyzed in a separate article.

Overall, this study describes the structural framework of AHN in terms of numbers of new neurons. Obviously, the numbers produced by our model might not be totally accurate, as there are certain caveats in the data used and further variability is expected between rodent strains and individuals. However, the predictions of the model agree well with expected levels of neurogenesis based on available empirical data and the general assumptions of the field, and therefore they might contribute to provide context to design experimental studies, to interpret findings and elaborate models of hippocampal function. In this regard, functional models of adult neurogenesis have traditionally acknowledged the possible impact of low levels of neurogenesis at older ages but have rarely included actual numbers in the discussion. Our study provides such estimations of neurogenesis across age that might contribute to produce more accurate and evidence-based functional models.

Finally, our interpretation of the results described here is that even if the low fraction of new neurons at the peak of neurogenesis showing preferential involvement in dentate function might have a relevant impact in the dentate network, their rapid, steep decrease during the first months of adulthood challenges that notion in middle aged and old individuals. Thus, we find very difficult to reconcile –from a computational and from a commonsense perspective-that such low number of new neurons might have such an essential role in the variety of functions in which they have been involved.

