## Supplementary data for "Meta-analysis: a quantitative model of adult neurogenesis in the rodent hippocampus"

For the model proposed in this manuscript, we searched for data combining BrdU injections with different survival time in rodents (both mice and rats) and cellular characterization of labeled cells using markers of differentiating cells such as DCX, PSA-NCAM and CR, and markers of mature cells such as NeuN and CB to identify functionally distinct new neurons 4-8 weeks old. Data for the model was obtained from the studies described below (Brandt et al., 2003; Brown et al., 2003; McDonald and Wojtowicz, 2005). Data was obtained from the text/tables when available or estimated from graphs using plotdigitizer software (available online at https://plotdigitizer.com).

**Brandt et al., 2003**

Six-week-old female C57BL/6 mice were injected with a single BrdU injection (50 mg/kg body weight) and were analyzed at 4 hours, 1 day, 3 days, 7 days, 18 days, and 28 days after the injection. 5 animals were analyzed at each timepoint except at 18 days (n=4).

Authors analyzed total number of cells co-labeled with BrdU and CR, as a marker of differentiating neurons, and BrdU and NeuN as marker of differentiated neurons.

The data from Brandt et al., 2003 lacks a day 7 timepoint on the expected peak of BrdU+/DCX+ cells. However, the overall curve is very similar to those including such timepoint and illustrate the similarities in the distribution of DCX+ cells along time between rats and mouse. They provide data on BrdU/NeuN labeled cells at 4 weeks, an early time point to assess the number of young, differentiated neurons, since some neurons may exhibit delayed or below detection NeuN expression. However, as reported in their Fig. 2, the population of BrdU/NeuN cells represents almost 100% of the cells labeled with BrdU at that time, indicating that all newly generated cells are accounted for (**Table S1**).

**Brown et al., 2003**

Eight-week-old female Wistar rats received intraperitoneal injections with BrdU (50 mg/kg body weight) at 2 hours and 4, 7, 10, 14, and 21 days, and 1, 2, 3, 4, 6, 9, 14, and 19 months timepoints. Each time point consisted of 4 animals. They did not report the total number of BrdU labeled cells at each timepoint and we used the data provided by MacDonald and Wojtowicz 2005 (**Table S1**).

**MacDonald and Wojtowicz 2005**

Young (38-day-old) and aging (12-month-old), male, Sprague–Dawley rats received two injections of BrdU (200 mg/kg each) 12 h apart, and were analyzed 1, 3, 7, 10, 14, 21, 28 and 60 days after the injections for proportion of BrdU, BrdU/DCX and BrdU/CB labeled cells. Three animals were studied at each timepoint (**Table S1**).

Other similar studies included (Kempermann et al., 2003) in mice, but used 12 consecutive injections of BrdU and therefore their data do not provide temporal resolution. Also, (Snyder et al., 2009) performed a similar longitudinal study in mice and rats, but they used animals injected with kainate to induce seizures, a confounding factor that might alter the molecular profile of labeled cells (Gruber et al., 1994; Maglóczky and Freund, 1995; Carter et al., 2008; Tallent and Qiu, 2008). They also reported non-trivial levels (up to 32 neurons/$mm^3$) of BrdU+/NeuN+ cells in the neocortex and striatum, an improbable finding that raise further questions about the reliability of the results in the dentate gyrus, and therefore we did not include their data in the analysis.

**Explanation of the model to quantify functional young new neurons 4-8 weeks old.**

In brief, the model requires figuring out the ratio between DCX expressing cells at any time and the number of new functional neurons 4-8 weeks old. We can use BrdU/DCX and BrdU/CR to identify and follow a cohort of new neurons along the differentiating phase, and BrdU/NeuN and BrdU/CB to identify and follow the same cohort when they become distinctly functional new neurons (DFNs).

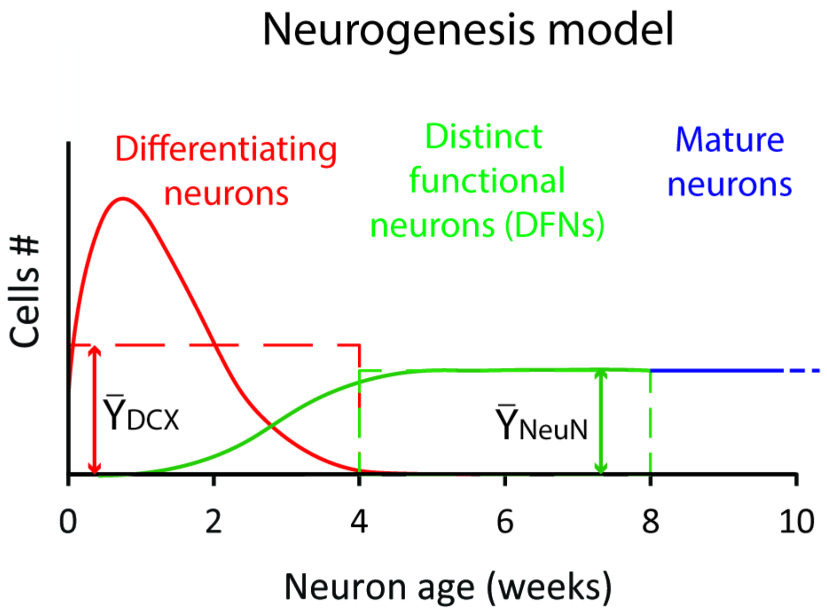


***Fig S1**. **Model of neurogenesis in the postnatal dentate gyrus**. A cohort of new neurons labeled with BrdU will have a ~4-week period of differentiation characterized by expression of (DCX, PSANCAM or CR; red curve) and a ~4-week period of maturation where it exhibits enhanced plasticity and excitability (DFNs) characterized by loss of differentiation markers and increased expression of maturation markers (NeuN and CB; green curve). After ~8 weeks, new neurons become physiologically indistinguishable from developmentally generated neurons and become*

*mature granule cells and continue expressing NeuN and CB. The average values of BrdU/DCX labeled cells ($\bar{Y}_{DCX}$) and the average values of BrdU/NeuN ($\bar{Y}_{NeuN}$) are represented with discontinuous lines in red and green respectively. Note that the area under the red curve equals the area of the polygon described by the average value of the distribution ($\bar{Y}_{DCX}$) between 0 and 4 weeks. Similarly, the area under the green curve (between 4 and 8 weeks) matches the area of the polygon whose height is the average of the distribution ($\bar{Y}_{NeuN}$) between 4 and 8 weeks. As the width of both polygons is the same (4 weeks), the ratio of both areas equals $\bar{Y}_{DCX} / \bar{Y}_{NeuN}$.*

The process of neurogenesis resembles the model in **Fig. S1**. There is an initial increase of BrdU/DCX labeled cells (red curve) that reaches a peak about 7 days after injection, likely reflecting progressive expression of DCX plus initial proliferation of DCX progenitors, followed by a marked decrease in the DCX/BrdU labeled population likely due to cell death and downregulation of DCX expression. Meanwhile, BrdU/NeuN cells (green curve) appear at around 1 week of age and increase their proportion to reach about 40% of the peak number of DCX+ cells around 4 weeks and become essentially stable thereafter. In this scenario, we need to relate the non-linear distribution of differentiating neurons in a single cohort of new neurons (red line in Fig S1) to the total population of DCX expressing cells. Since DCX expression lasts for about 4 weeks, the total number of DCX labeled cells at any point will pool all cohorts of differentiating neurons generated during those 28 days.

Mathematically, the total number of DCX labeled cells will be the integral of the function describing their distribution along time (4 weeks) multiplied by the number of cohorts produced in those 4 weeks, that is unknown. Similarly, the total number of DFNs will be the integral of the function describing those new neurons along time (4 weeks, from week 4 to week 8) multiplied by the number of cohorts. Because we are interested in the ratio between those two factors, the number of cohorts can be canceled, and the ratio we are looking for becomes the ratio between both integrals.

We could attempt to find the equations fitting the curves and proceed to integrate. However, we can use a geometrical shortcut. The integral of a function corresponds to the area bound by the function and the x axis. Thus, the integral of differentiating neurons corresponds to the area bound by the function (red curve) in Fig S1 between 0 and 4 weeks, that is equivalent to the area of a polygon whose height is the average value of the distribution between 0 and 4 weeks ($\bar{Y}_{DCX}$). Similarly, the integral of the function describing the population of DFNs equals the area of the polygon whose height is the average value of the distribution between 4 and 8 weeks ($\bar{Y}_{NeuN}$). Furthermore, since the width of both of those polygons is the same (4 weeks), the ratio can be simplified to the quotient of the average (height) of both distributions ($\bar{Y}_{DCX} / \bar{Y}_{NeuN}$).

To calculate the average of the number of differentiating new neurons ($\bar{Y}_{DCX}$), we interpolated daily values for the differentiating cells (BrdU and DCX/CR labeled; **Table S1, S2**). For DFNs, the function is linear, and we calculated the average between the values at 28 and 60 days (**Table S1, S2**). The resulting averages were 41% (of the maximum) for differentiating neurons and 37% of the maximum for DFNs (**Table S2** and **Fig 2B** in main text), that rendered a ratio of

0.907 or 91%. Thus, the population of new functional neurons 4-8 weeks old is about 91% of the total population of DCX expressing cells present on average 4 weeks before as shown in **Fig 2B**.

To obtain an equation describing the number of DFNs, we transformed the distribution of DCX labeled cells provided by equations 1 and 2 (in the main text) by multiplying each value by 0.91 and assigning them a 4-week delay. Regression of the resulting data produced equations 3 and 4 (in the main text. Another possible approach would be to perform regression of the transformed (*0.91 and added 4-week delay) raw data points of DCX labeled cells, that rendered very similar equations and total number of generated neurons. For example, for the rat, the equation produced by the latter procedure was $y= 149254x^{-1.467}$ and the total number of DFNs along the lifespan was 114,220, both very similar to the values obtained with the first procedure described in the main text.

**Limitations of the model**

It can be argued that one caveat of the model is the heterogeneity of the data used to build it, as compiles data from different markers and time points of analysis. Although this variability warrants caution when interpreting the results, it is noteworthy that the predictions of the model are in tune with the levels of neurogenesis expected from the distribution of DCX labeled cells and also agree with predictions based on experimental data (Ninkovic et al., 2007; Imayoshi et al., 2008; Encinas et al., 2011; Lazic, 2012; Pilz et al., 2018). The data suggest similar parameters of neurogenesis between mice and rats, although with slightly lower levels of neuronal survival in mice. In this regard, (Snyder et al., 2009) reported large differences in neuron survival between rats and mice, but as described above, their data might not be totally reliable. Regarding inter-strain variations, (Kempermann et al., 1997) reported that C57BL/6 animals exhibited higher levels of neurogenesis compared to other common strains such as CD1 and BALB/c, and more than double number of putative new neurons than 129/SvJ animals. Therefore, considering our model is based on C57BL/6 animals, it could be argued that we are likely overestimating the level of neurogenesis in other strains of mice. Nonetheless, we found that the normalized distributions of differentiating new neurons and DFNs were quite similar across species, strains, and labs, as reflected in **Fig. 2A** and therefore we combined them to obtain an average value.

Another issue is the age of the animals whose data were used for our model. Most were 5-8 weeks old when injected with BrdU, meaning they were adolescent and not adult, as discussed in the main text and extensively reviewed elsewhere (Arellano et al., 2024). This might imply that there could be differences in cell differentiation dynamics and survival ratios that might change in adult and aging animals. However, most studies have pointed no differences in such parameters (McDonald and Wojtowicz, 2005; Rao et al., 2005, 2006; Hattiangady and Shetty, 2008) and the datasets in our model comparing very young (adolescent) P38 with aged 12-month-old animals only showed differences in the rate of new neuron survival, that was slightly lower in older animals. An alternative view of this potential caveat is that the model includes heterogeneous data that might overall produce a more averaged model of the neurogenic process regardless of age, species, or strain. Indeed, the purpose of this model is not to produce a precise and accurate prediction of the number of new neurons, but an approximate estimate of the number of new neurons that makes sense conceptually and agrees with experimental studies and with empirical data such as the number of differentiating new neurons labeled with DCX. The main purpose of the model is provide rough estimates to contextualize the potential functional impact of adult neurogenesis in realistic quantitative terms, and offering perspective regarding the limitations imposed by very low numbers of new neurons produced in aged animals, as most functional hypotheses about this trait have been derived from studies in very young animals with relatively high levels of neurogenesis, disregarding the steep decline of neurogenesis in middle aged and old animals.

**Note on Cole et al., 2020**

In figure 8A, (Cole et al., 2020) provide an equation that describes the number of new neurons produced along the lifespan of a rat. However, we noticed a typo in the equation: the first constant, 1213 should be 12130.
